# Condensate formation of the human RNA-binding protein SMAUG1 is controlled by its intrinsically disordered regions and interactions with 14-3-3 proteins

**DOI:** 10.1101/2023.02.09.527857

**Authors:** John Fehilly, Olivia Carey, Eoghan Thomas O’Leary, Stephen O’Shea, Klaudia Juda, Rahel Fitzel, Pooja Selvaraj, Andrew J. Lindsay, Bálint Mészáros, Kellie Dean

## Abstract

SMAUG1 is a human RNA-binding protein that is known to be dysregulated in a wide range of diseases. It is evolutionarily conserved and has been shown to form condensates containing translationally repressed RNAs. This indicates that condensation is central to proper SMAUG1 function; however, the factors governing condensation are largely unknown. In this work, we show that SMAUG1 drives the formation of liquid-like condensates in cells through its non-conventional C-terminal prion-like disordered region. We use biochemical assays to show that this liquid-liquid phase separation is independent of RNA binding and does not depend on other large, disordered regions that potentially harbor several binding sites for partner proteins. Using a combination of computational predictions, structural modeling, *in vitro* and in cell measurements, we also show that SMAUG1-driven condensation is negatively regulated by direct interactions with members of the 14-3-3 protein family. These interactions are mediated by four distinct phospho-regulated short linear motifs embedded in the disordered regions of SMAUG1, working synergistically. Interactions between SMAUG1 and 14-3-3 proteins drive the dissolution of condensates, alter the dynamics of the condensed state, and are likely to be intertwined with currently unknown regulatory mechanisms. Our results provide information on how SMAUG1 phase separation is regulated and the first known instance of 14-3-3 proteins being able to completely dissolve condensates by directly interacting with a phase separation driver, which might be a general mechanism in cells to regulate biological condensation.

**Graphical abstract:** 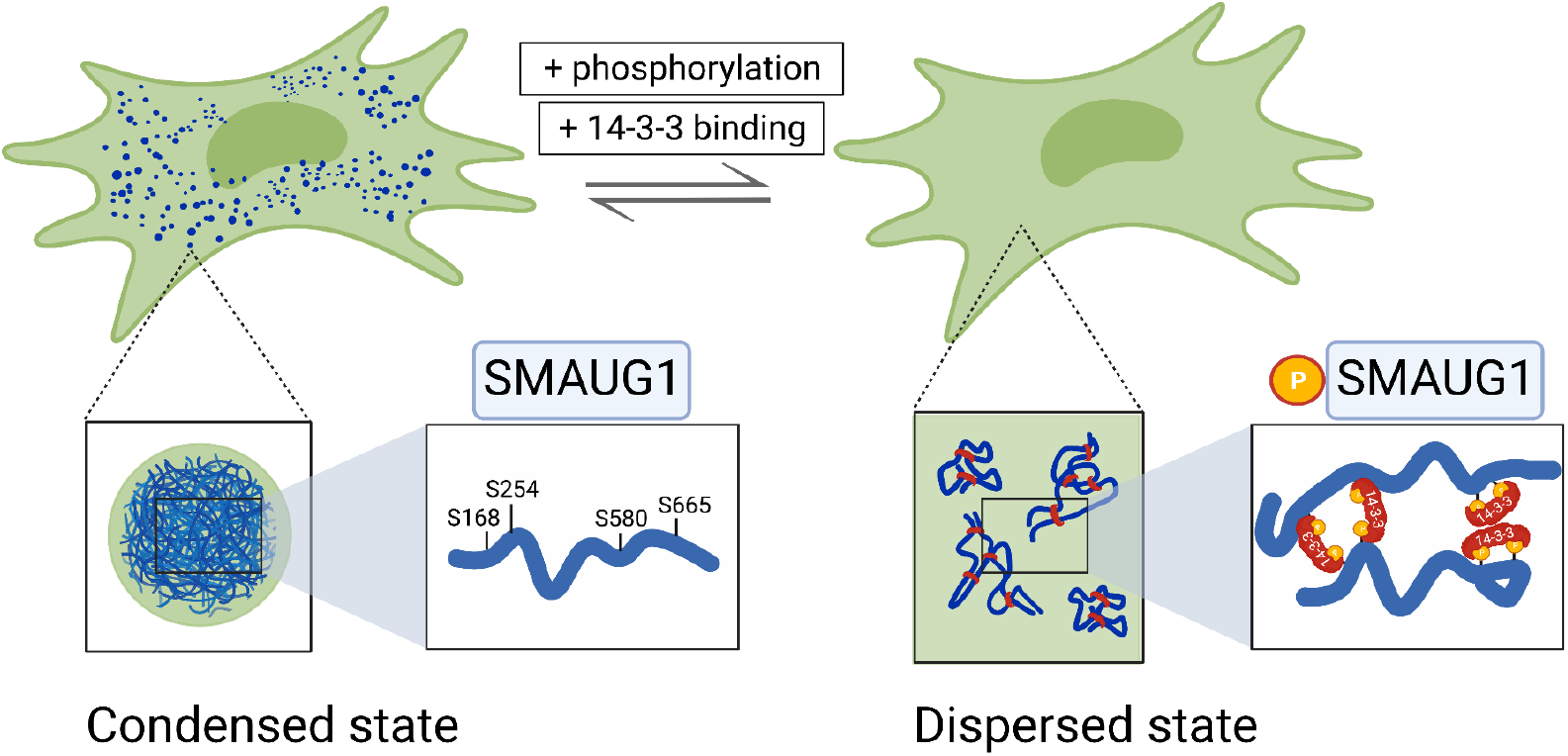

**Highlights:**

- SMAUG1 is a human RNA-binding protein capable of condensation with unknown regulation
- A prion-like domain of SMAUG1 drives condensation via liquid-liquid phase separation
- SMAUG1 interacts with 14-3-3 proteins via four phospho-regulated short linear motifs
- 14-3-3 interactions change the dynamics of SMAUG1 condensates, promoting their dissolution
- This is the first described regulatory mechanism for SMAUG1-driven condensation

## Introduction

SMAUG1 (SAMD4A) is a 718 residue, human RNA-binding protein that is highly evolutionarily conserved from yeast to humans, functioning as a translational repressor. Diverse SMAUG1 functions have been explored in multiple organisms: in *Drosophila melanogaster*, SMAUG1 directs body axis formation through its repression of *nanos* mRNA [1,2], is involved with global regulation of thousands of mRNAs in the early fly embryo [3], is essential for maternal-to-zygotic transition [4], and has been shown to specifically destabilize mRNAs encoding mitochondrial proteins leading to mitochondrial dysfunction [5]. In mice, SMAUG1 has been linked to bone formation [6] and energy production, with mutations in the protein leading to uncoupled mitochondrial function [7]. Human SMAUG1 has also been shown to affect mitochondrial functions [8], as well as being linked to a range of diseases, including muscular dystrophy (myotonic dystrophy type 1 (DM1) [9] and oculopharyngeal muscular dystrophy [5]), as well as suppression of the hepatitis B virus [10].

This diverse set of functions is reflected in the complex architecture of SMAUG1. RNA-binding is directed by the sterile alpha motif (SAM) [11], which has a highly conserved sequence. The rest of the protein largely lacks a stable structure, being composed of multiple intrinsically disordered regions (IDRs) according to AlphaFold2 structural models [12,13]. While the sequences of these large IDRs are poorly conserved, they are present in every known homolog of SMAUG1, indicative of a functional role. We have virtually no mechanistic knowledge about the functions of these IDRs; however, based on the general functional repertoire of IDRs [14,15], it is plausible that the protein-protein interactions formed by SMAUG1, as well as various regulatory functions, are likely mediated here.

Recent works have shown that a major function of IDRs, especially low complexity ones, is to drive biomolecular condensation of proteins, often via liquid-liquid phase separation. Condensation properties of SMAUG1 have been observed across a wide evolutionary range. The yeast (*S. cerevisiae)* homologue of SMAUG1, called Vts1, assembles into gel-like condensates with poor dynamic properties, promoted by an IDR [16]. In fly, Smaug forms cytosolic bodies [17], and in human neurons SMAUG1 forms silencing foci (S-foci) that are thought to contain translationally repressed mRNAs [18,19]. Biological condensation has been recognized as a major regulatory mechanism for proteins involved in a wide range of biological processes, including modulation of chemical reaction rates, governing molecular availability, structural organization of the cell and regulating homeostasis [20]. Low complexity IDRs have been shown to drive liquid-liquid phase separation [21], creating condensates that are linked to healthy cell functions as well as disease states [22–24]. These works have also shown that since condensation has critical cellular roles, it is very tightly regulated through protein abundance, interactions with proteins, RNAs and other macromolecules, as well as through post-translational modifications, most often phosphorylation.

While SMAUG1 performs important functions and participates in biological condensation in cells, it is unclear how its function and its phase separation propensity are regulated. Earlier observations in fly show that phosphorylation of Smaug by Fused (downstream of Smoothened, a G-protein coupled receptor (GPCR) involved in hedgehog signaling) reduces its ability to form cytosolic bodies [17]. In human U-2 OS cells, SMAUG1 foci are reported to dissolve in response to electron transport chain complex 1 inhibition and AMPK activation/mTOR inhibition [8]. Upon specific synaptic stimulation through the N-methyl-D-aspartic acid (NMDA) receptor, the S-foci dissipate and presumably release mRNAs that can be translated, including the release of Ca^+2^/calmodulin-dependent protein kinase II (CaMKIIa) mRNA [19]. While this information clearly shows that phosphorylation and interactions with proteins are crucial in SMAUG1 regulation, they do not provide information about which residues in SMAUG1 are involved, hindering mechanistic understanding of SMAUG1 regulation and function.

In this work we set out to identify regions, modifications, and interactions of human SMAUG1 that govern its ability to participate in or drive biomolecular condensation. We use computational predictions to guide our experimental work with tagged-versions of wild-type and mutant versions of SMAUG1 in cells to explore the following questions: Which region(s) of SMAUG1 drives phase separation; is SMAUG1-driven phase separation RNA-dependent; which SMAUG1 regions form protein-protein interactions that modulate the phase separation capacity and dynamics of the resulting condensates; and how are these interactions regulated through phosphorylation.

## Materials and Methods

### Cell lines and cell culture

U-2 OS osteosarcoma cells were purchased from the American Tissue Type Collection (ATCC) (Manassas, VA, USA). Human embryonic kidney cells (HEK293) were a kind gift from Prof Mary McCaffrey (UCC, Cork, Ireland). Cell lines were authenticated using short tandem repeat (STR) profiling (Eurofins Genomics). Cells were cultured according to the following media requirements. U-2 OS cells were maintained in McCoy’s 5A media (Merck) supplemented 10% fetal bovine serum (FBS) and 1% penicillin/streptomycin. HEK293 cells were maintained in Dulbecco’s Modified Eagle’s Medium (DMEM) (Merck) supplemented with 10% FBS and 1% penicillin/streptomycin. Cells were maintained at 37°C with 5% CO_2_ and were mycoplasma-free.

### Modification of DNA vectors for mammalian cell expression

For transient expression of tagged versions of SMAUG1 and 14-3-3 proteins, we created two vectors by modifying pCMV (gift from Dr Paul Young, UCC, Cork, Ireland). First a His_6_-TEV-2xFLAG tag was synthesized as a gene block (Integrated DNA Technologies, IDT), and the fragment was ligated into the *BamHI/XbaI* sites of pCMV. Second, a Myc tag was generated by annealing two DNA oligonucleotides that had *BamHI* and *XbaI* overhangs. A *SpeI* site was introduced to facilitate selection of positive clones. To do this, oligonucleotides that encode the Myc tag were synthesized (IDT) and mixed in equal molar amounts. 10 μl of 10x annealing buffer (400 mM Tris/HCl pH7.9, 500 mM NaCl, and 100 mM MgCl2) and sterile distilled water was added to bring the final volume to 100 μl. This was then incubated at 95°C for 10 minutes and slowly cooled to room temperature. The annealed oligos were phosphorylated using T4 polynucleotide kinase (PNK, New England Biolabs). The phosphorylated and annealed oligos were then diluted in Tris-EDTA (TE) buffer before ligation into *BamHI/XhoI* digested pCMV vector.

### Cloning and mammalian cell expression constructs for SMAUG1 and 14-3-3 proteins

Full-length human SMAUG1 corresponds to the protein sequence in UniProt under the accession Q9UPU9 (SMAG1_HUMAN) referring to the canonical isoform of 718 residues. *SMAUG1* coding sequence (CDS) (NCBI RefSeq NM_015589.6) was amplified by polymerase chain reaction (PCR) using complementary DNA oligonucleotide primers from a human skeletal muscle cDNA library (gift from Dr Paul Young, UCC, Cork, Ireland) and cloned into pCMV-HisTEV2XFLAG vector. Full-length 14-3-3 human proteins, with associated UniProt accession identifiers, were used: 14-3-3β/a P31946 (1433B_HUMAN); 14-3-3γ P61981 (1433G_HUMAN), 14-3-3η Q04917 (1433F_HUMAN) and 14-3-3ζ/d P63104 (1433Z_HUMAN). To make expression constructs, the corresponding CDS from *YWHAB* (NCBI RefSeq NM_003404), *YWHAG* (NCBI RefSeq NM_012479), *YWHAH* (NCBI RefSeq NM_003405) and *YWHAZ* (NCBI RefSeq NM_001135699) were amplified by PCR using complementary DNA primers from a human skeletal muscle cDNA library as before and cloned into pCMV-myc vector. 14-3-3γ CDS was amplified by PCR and subcloned into pCMV-mCherry (Clontech). Deletion and point mutations of SMAUG1 were created using Q5 site-directed mutagenesis kit (NEB). Mutagenic primers were designed using the NEBaseChanger tool and synthesized (IDT). PCR conditions were followed according to each primer pair as recommended by NEB using pCMV-SMAUG1-HisTEV2xFLAG as the template. Each deletion construct of SMAUG1 refers to the protein sequence with the region removed defined after the name of the construct as shown in Figure 2A: ΔIDR:237-311, ΔSAM:321-381 and ΔPLD:567-605. Serine-to-alanine mutations were introduced at amino acid positions, 168, 254, 580 and 665 in SMAUG1. The coding sequences of wild-type SMAUG1 and S168/254/580/665A (quad S/A mutant), with disabled 14-3-3 binding motifs, were subcloned in frame from their corresponding SMAUG1-HisTEV2xFLAG plasmids into pEGFP-N1 (Clontech) using *XhoI* and *BamHI*, to create C-terminal, GFP-tagged versions of wild-type and mutant SMAUG1. All constructs were verified by Sanger sequencing (Eurofins Genomics), with alignments performed in Benchling. The DNA sequences of all gene blocks and oligonucleotides that were used for vector construction, PCR amplification, mutagenesis and Sanger sequencing are listed in Supplementary Table S1.

**Figure 1:**
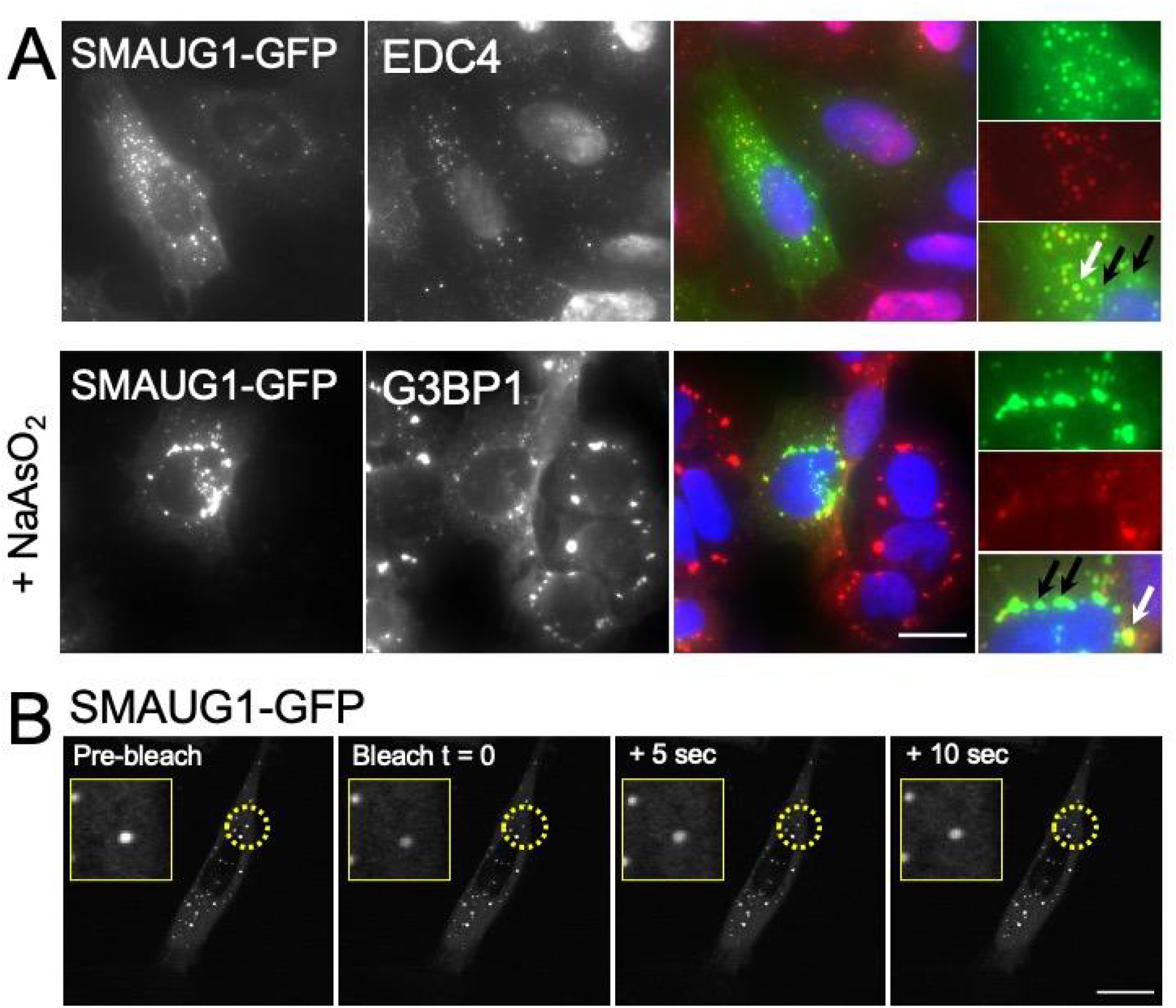
SMAUG1 forms condensates with liquid-like properties that are distinct from P-bodies and stress granules. (A) Immunofluorescence of EDC4 and G3BP1 in U-2 OS cells expressing SMAUG1-GFP. To induce SG formation, cells were treated with sodium arsenite (10μM) for 1 hour. White arrows indicate co-localization of SMAUG1 with P-body and SG markers; black arrows point to distinct SMAUG1 condensates. Images were collected at 60X magnification. (B) SMAUG1 shows rapid recovery in FRAP. U-2 OS cells transiently expressing SMAUG1-GFP were seeded onto glass bottomed plates and analyzed 24 hours post-transfection. Dashed circles surround a condensate that shows a reduction in intensity upon bleaching (zoomed images pre- vs. bleach, time = 0), followed by fast recovery at 5- and 10-seconds post-bleach. All images are representative of experiments from three biological replicates. Scale bar is 20μm. For videos of FRAP of SMAUG1-GFP see Supplementary video 1.

### Immunofluorescence of processing bodies and stress granules

SMAUG1-GFP was transfected into U-2 OS cells via electroporation using a Lonza Nucleofector 2B (program X-001). Following electroporation, cells were transferred to a culture dish, containing glass coverslips. To induce stress granule formation, cells at 24-hours post-transfection were treated with sodium arsenite (Sigma Aldrich) at a final concentration of 10mM for 1 hour. Cells were fixed with 4% paraformaldehyde and incubated with permeabilization/blocking buffer (5% normal goat serum, 2% bovine serum albumin, 0.2% Triton X-100) for 30 mins at room temperature. Cells were incubated with primary antibodies (diluted 1:50) directed against EDC4 (Santa Cruz Biotechnology, SC-8418, mouse) and G3BP1 (Bethyl Laboratories, A302-033A, rabbit) in antibody buffer (5% normal goat serum, 2% bovine serum albumin) overnight at 4°C. Coverslips were washed three times with phosphate-buffered saline (PBS) and then incubated with anti-mouse or anti-rabbit Cy3 conjugated secondary antibodies (Jackson ImmunoResearch, dilution 1:1000) in antibody buffer for one hour at room temperature. Cell nuclei were stained with 4’,6-diamidino-2-phenylindole (DAPI). Coverslips were then washed three times with PBS and mounted onto glass slides. Cells were imaged using an EVOS FL Auto Imaging System (Thermo Fisher) using the 60x objective. Images were processed using Fiji [25]. For each condition 5 images were taken (approximately 15 cells/image) per condition across three biological replicates.

### Fluorescence recovery after photobleaching (FRAP)

The FRAP experiments were performed using a Zeiss LSM 510 Meta confocal microscope with a 63X 1.4 NA Plan Apo oil immersion objective. SMAUG1-GFP constructs (wild-type and mutant) (3ug) were transfected into U-2 OS cells via electroporation as before. Cells were plated onto 35mm glass-bottomed dishes (MatTek). The cells were imaged approximately 24-hours post-transfection. To perform FRAP, a single region of interest (ROI) in each cell was bleached by eight iterations at 100% power with the 488 nm argon laser. Images were recorded every 1 second for approximately 2 minutes post-bleach. The average fluorescence intensity in the ROI was calculated at each time point and was normalized against the average fluorescence intensity of the whole cell at the same time point using the FRAP Profiler plugin in ImageJ. The mobile fraction (MF) was calculated using the formula MF = ((ymax - ymin)/(1 - ymin))*100. Only ROIs in which the fluorescence intensity was bleached by greater than 70% were analyzed. For the complete FRAP data, see Supplementary table S2; for FRAP videos of wild type and quad S/A mutant GFP-tagged SMAUG1 see Supplementary videos 1 and 2.

### Disorder and prion-like propensity predictions

The disordered tendency of SMAUG1 was assessed using IUPred2A [26], which returns two values for each residue; one for the probability of that residue having no fixed structure (IUPred2) and the probability of that residue lacking structure and being able to interact with a folded partner protein (ANCHOR2) (Figure 1B). These scores were generated using the online version of IUPred2A using default settings (long disorder prediction). Prion-like propensities were assessed using PLAAC [27] setting the background to 100% human.

### Endogenous 14-3-3 co-immunoprecipitation with SMAUG1-GFP

HEK293 cells were seeded in 15cm plates 24 hours prior to transfection. Cells were transfected at 70% confluency using calcium phosphate transfection. Cells were mock transfected (identical transfection reagents with no DNA) or transfected with 22.5μg pEGFP-N1 or SMAUG1-GFP. Cells were harvested 48 hours post transfection. Immunoprecipitations were carried out using GFP-Trap agarose (Chromotek) according to the manufacturer’s instructions. Cells were washed three times in ice-cold PBS and scraped from the plates. Cells were lysed in lysis buffer containing 10 mM Tris/HCl, pH 7.5, 150 mM NaCl, 0.5 mM ethylenediaminetetraacetic acid (EDTA), 0.5% Nonidet P40 substitute (Sigma Aldrich) supplemented with cOmplete protease inhibitor cocktail (Roche) and Halt phosphatase inhibitor cocktail (Thermo Fisher). Lysates were incubated by tumbling end-over-end with 25μl of bead slurry at 4°C for 1.5 hours. Beads were washed three times with a wash buffer identical to the lysis buffer, but without NP40 substitute. To elute bound proteins from the beads there were incubated with 2x sodium dodecyl sulfate (SDS) sample buffer (120 mM Tris/Cl pH 6.8; 20% glycerol, 4% SDS; 0.04% bromophenol blue; 10% β-mercaptoethanol) at 95°C for 10 min, followed by SDS-PAGE and immunoblotting (see below). 4% of the total lysate was loaded into the input and unbound lanes and 50% of the total immunoprecipitated material was loaded into the bound lane.

### SMAUG1/14-3-3-myc co-immunoprecipitations with RNase A treatment

HEK293 cells were transfected with 14-3-3-myc tagged constructs, and cell lysate was prepared as previously described. Co-immunoprecipitation using Myc-Trap beads (Chromotek) was performed as described above for the GFP-Trap beads. Prior to the wash stage after incubation of beads with cell lysate, the bead/cell lysate sample was split into two pre-chilled 1.5 ml tubes. One sample was treated with RNase A. Lysate and beads were incubated with 50 μl of 10 mg/ml RNaseA at 37°C for 10 minutes. The other sample was stored on ice. After RNase A treatment, the beads were magnetically separated and 50 μl of the unbound sample was stored for immunoblot analysis (input), with the remaining supernatant being discarded. Beads were resuspended and washed in dilution buffer, before once again being separated magnetically. The supernatant was then discarded, and the beads were washed twice more in dilution buffer. After washing the beads were resuspended in 100 μl of 2x SDS sample buffer. Resuspended beads were boiled at 95°C for 10 minutes to dissociate the immunocomplexes. The beads were then magnetically separated, and SDS-PAGE and immunoblot analysis (see below) was performed with 30 μl of the protein sample.

### Generating and evaluating complex structural models with FoldX and FlexPepDock

FoldX [28] version 4 was used to generate structural models of various peptides derived from human SMAUG1 bound to human 14-3-3γ. As a template, the PDB structure 6a5s containing a peptide centered on the phosphorylated Ser211 of transcription factor EB (TFEB) bound to human 14-3-3γ [29] was used. Only the 3 residues preceding and following the pSer residue were kept. The structure was first relaxed with the RepairPDB function. Next, the ±3 residues around the pSer were changed to match the sequence of the SMAUG1 peptide being evaluated using the BuildModel command, defining the required sequence changes as mutations. The generated models were used as inputs for backbone optimization with FlexPepDock [30,31]. 300 high-resolution structures were generated and the highest-scoring predicted complex structure was optimized for side-chain conformations with FoldX using the repairPDB command, yielding a predicted ΔG value for the complex between the domain and the peptide (ΔG_complex_). In addition, the structure was also repaired after deleting the peptide atoms, leaving the monomeric domain structure, yielding ΔG_domain_ estimating the stability of the domain. From this, the predicted free energy of the peptide binding was calculated as ΔG_binding_=ΔG_complex_-ΔG_domain_.

### Phosphorylation sites

Phosphorylation data for human SMAUG1 was taken from PhosphoSitePlus [32] version 6.6.0.2. Both low- and high-throughput phosphorylations were used.

### Assessing structural accessibility using AlphaFold2

The AlphaFold2 computed structural model of human SMAUG1 was taken from version 4 of the AlphaFoldDB [13]. The relative solvent accessible surface of each residue was calculated by running DSSP on the AlphaFold structure and comparing the calculated solvent accessible surface area in the structure to that calculated in a GGXGG conformation. This relative accessibility was averaged in a ±20 residue window, as this measure was found to correlate well with the presence of high flexibility/disorder in benchmarks [12]. Regions of residues with relative accessibility values above 0.55 were considered as suitable candidates for harboring true 14-3-3 binding motifs.

### Predicting 14-3-3 motifs from the sequence

Candidate 14-3-3 binding motifs were predicted in the SMAUG1 sequence using the Eukaryotic Linear Motif resource [33] and the 14-3-3-Pred method [34].

### Phostag SDS-PAGE to determine SMAUG1 phosphorylation status

HEK293 cells were transfected with pCMV-SMAUG1HisTev2xFlag vector to allow for the transient overexpression of SMAUG1. Cells were harvested two days post-transfection using a NP-40 lysis buffer (50mM Tris-HCl pH 8, 150mM NaCl, 1% NP-40, supplemented with c0mplete tablet mini-EDTA-free protease inhibitor (Roche) and Halt phosphatase inhibitor cocktail (Thermo Fisher Scientific)). Total protein concentration was quantified using a bicinchoninic acid (BCA) assay. Lambda protein phosphatase treatment was applied to lysates to compare phosphorylation of SMAUG1 in treated vs untreated samples. The lambda protein phosphatase procedure was carried out following the manufacturer’s guidelines (New England Biolabs). The sample was incubated at 30°C for 2 hours, and the phosphatase was heat-inactivated at 65°C for 1 hour. Lambda protein phosphatase treated and untreated samples were subjected to sodium dodecyl-sulfate polyacrylamide gel electrophoresis (SDS-PAGE) or Phos-tag SDS-PAGE for 2 hours. The SDS-polyacrylamide gels consisted of an 8% acrylamide separation gel and a 4% stacking gel. The Phos-tag gels consisted of a separating gel copolymerized with Phos-tag (8% acrylamide, 25μM acrylamide-pendant Mn^2+^ Phos-tag, 40μM MnCl_2_). After electrophoresis, Phos-tag gels were soaked in a transfer buffer (25mM Tris, 192mM glycine, and 20% methanol) containing 1 mM EDTA for 20 min with gentle agitation for elimination of the manganese ions from the gel. Next, the gels were soaked in a transfer buffer without EDTA for 10 min with gentle agitation. The resolved proteins were transferred to nitrocellulose membranes for immunoblotting (see below).

### Mutant SMAUG1 co-immunoprecipitations with 14-3-3γ myc

Mutant SMAUG1 co-immunoprecipitations with myc-tagged 14-3-3γ were carried out as described above with a slight modification. HEK293 cells were seeded in 10cm plates and transfected with 2.5μg 14-3-3γ-myc and 7.5μg of either WT SMAUG1-FLAG or various mutant SMAUG1-FLAG constructs (Ser to Ala mutants: S168A, S254A, S580A, S665A; S168/254A, S254/580A, S254/665A, S580/665A; S168/254/580A, S254/580/665A; or S168/254/580/665A). Immunoprecipitations were carried out using Myc-Trap magnetic agarose (Chromotek) according to the manufacturer’s instructions at 4°C for 2 hours. 3% of total material was loaded into the bound/unbound lanes and bound lanes were loaded with 25% of immunoprecipitated material.

### SDS-PAGE and immunoblotting

Sodium dodecyl sulfate-polyacrylamide gel electrophoresis (SDS-PAGE), followed by immunoblotting, was carried out using either 10 or 12% acrylamide gels. For all blots, PageRuler prestained protein ladder (Thermo Fisher) was used. Proteins were transferred to PVDF or nitrocellulose membranes and blocked for one hour with 5% nonfat milk or 2% bovine serum albumin (BSA) if probing for phosphoproteins. Membranes were usually incubated with the primary antibody at 4°C overnight, except for anti-b-actin, which was incubated at 4°C for two hours. The following primary antibodies and dilutions were used in our study: anti-SMAUG1 (Proteintech, 17387-1-AP, 1:1000), anti-AKT (Cell Signaling Technology, 2920, 1:1000) and anti-phospho-AKT Ser^473^ (Cell Signaling Technology, 4060,1:1000); anti-GFP (Proteintech, 50430-2-AP, 1:4000), anti-pan-14-3-3 (B-8) (Santa Cruz Biotechnology, sc-133233, 1:1000), anti-14-3-3η (D23B7) (Cell Signaling Technologies, 5521, 1:1000), anti-14-3-3ζ (Novus Biologicals, NBP2-24593SS, 1:1000). Following primary antibody incubation, membranes were washed three times in Tris-buffered saline with Tween 20 (TBST) (137mM NaCl, 2.7mM KCl, 19mM Tris base, 0.1% Tween) and were detected with secondary antibodies. For some blots, we used horseradish peroxidase (HRP)-conjugated isotype-specific anti-rabbit IgG or anti-mouse IgG (1:10000; DAKO, Cambridgeshire, UK) secondary antibodies, followed by enhanced chemiluminescence (ECL) (Pierce) (Figures 3B, 3C, 3D and 5A). For other blots, we used LI-COR anti-rabbit IRDye 800CW or anti-mouse IRDye 680RD (1:15000) secondary antibodies (Figures 3A, 5B and 5C), followed by imaging on an Odyssey imaging system (LI-COR).

**Figure 2:**
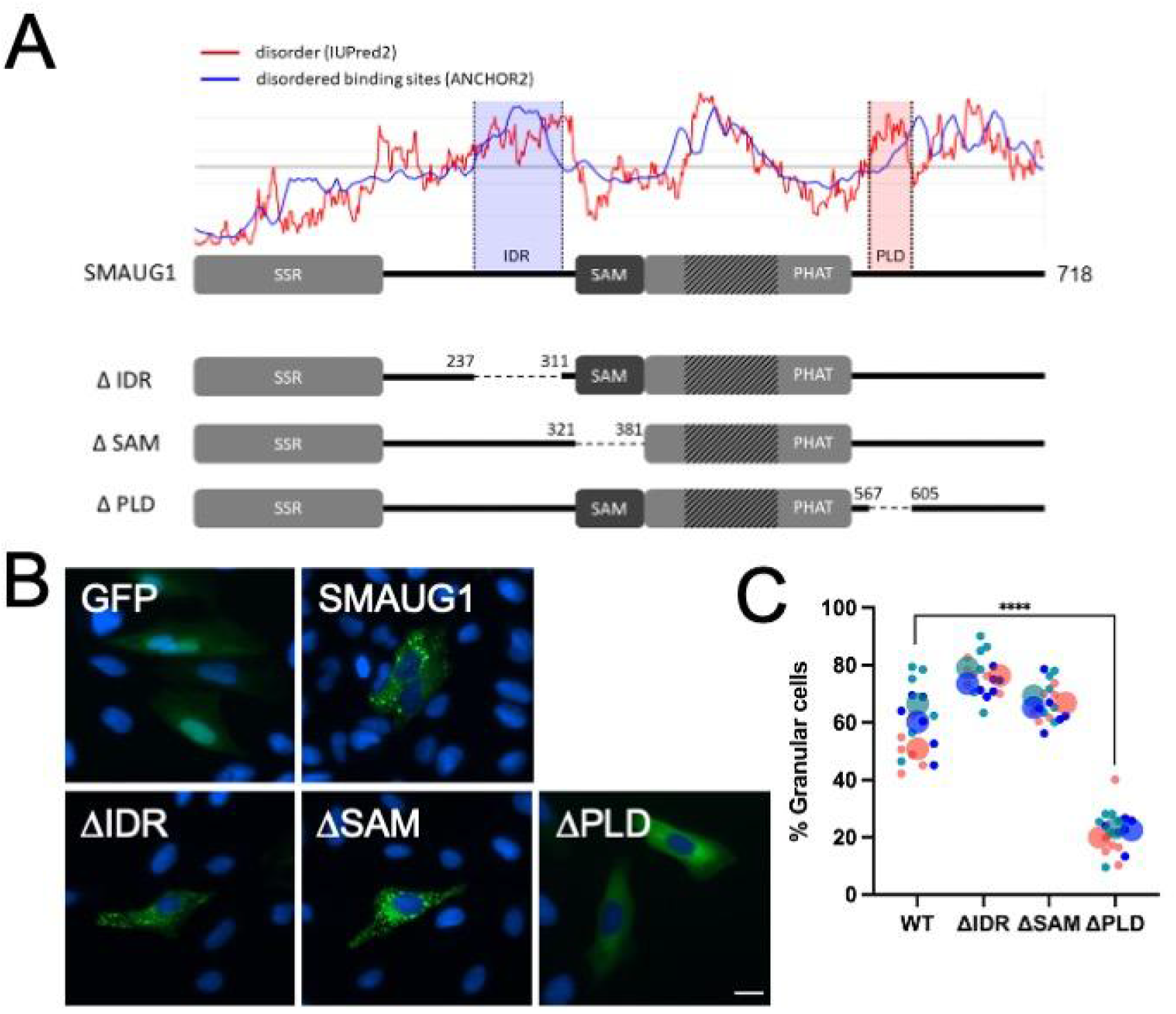
SMAUG1 is largely disordered, and condensate formation is driven by a low-complexity, prion-like domain. (A) Human SMAUG1 constructs used in this study. The red line in the top graph shows the predicted disorder tendency with residues likely to lack an intrinsic 3D structure having scores above the 0.5 line (IUPred2). The blue line indicates disordered binding regions with residues having scores above 0.5 being likely to be part of regions lacking 3D structure but binding to a structured protein partner (ANCHOR2). IDR - intrinsically disordered region, PLD - prion-like domain. In the domain diagrams below the prediction output, the following regions are marked: SSR - N-terminal SMAUG similarity region, SAM - sterile alpha motif (the RNA-binding region), PHAT - pseudo-HEAT analogous topology domain, a SMAUG-specific structural domain with a flexible insertion in human SMAUG1 is marked by shading in the figure. (B) EGFP-tagged deletion mutants of SMAUG1 were transiently expressed in U-2 OS cells. Representative images from three biological replicates are shown. (C) Superplots [44] graph showing the percentage of granular cells when transfected with WT SMAUG1 and deletion constructs. Each color represents a biological replicate; the large circle is the average % granular cells of the technical replicates (small circles; n=5). Means of each group were compared with a one-way ANOVA using GraphPad Prism 9 using the Tukey’s test to correct for multiple comparisons, p<0.001 (****).

**Figure 3:**
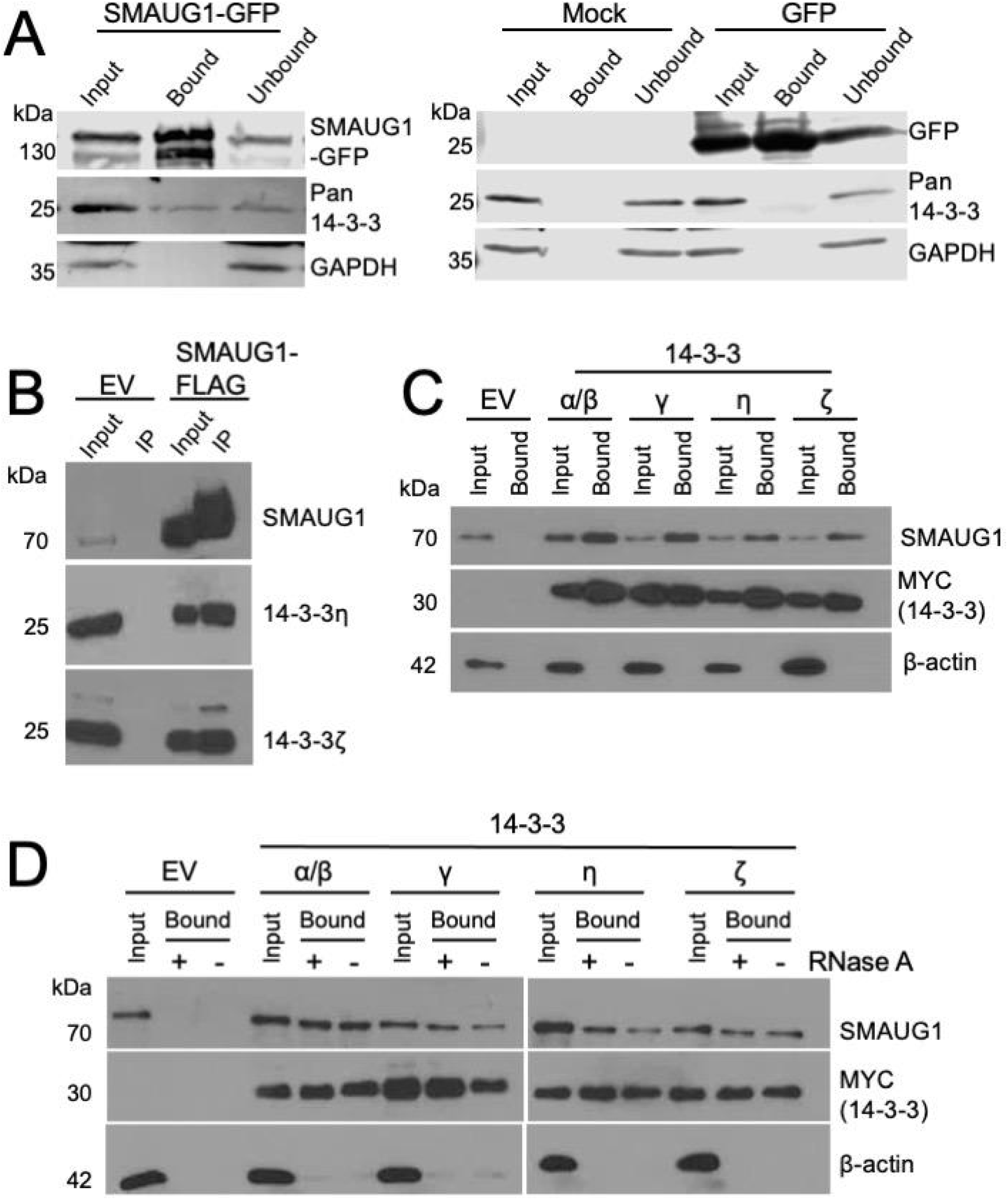
SMAUG1 interacts with members of the 14-3-3 protein family. (A) GFP-tagged SMAUG1 pulls down 14-3-3 proteins from HEK293 cell lysates. Left: identification of 14-3-3 proteins with a pan-14-3-3 antibody recognizing all seven members of the human 14-3-3 protein family (α/β, ε, γ, η, τ, ζ and σ), with GAPDH as the loading control. Right: controls using mock transfected, and cells transfected with GFP alone showing no interaction with any 14-3-3 proteins. (B) FLAG-tagged SMAUG1 pulls down both 14-3-3η and 14-3-3ζ. (C) Myc-tagged 14-3-3β, γ, η, and ζ proteins pull down endogenous SMAUG1 (14-3-3α denotes the phosphorylated version of 14-3-3β, and assuming that the interaction is mediated by a canonical 14-3-3/motif binding, the phosphorylation has no effect on the interaction). (D) Similar to panel (C), myc-tagged 14-3-3 pulls down endogenous SMAUG1 even after RNase treatment, showing that the interaction is independent of having intact RNA in the cell lysate.

**Figure 4:**
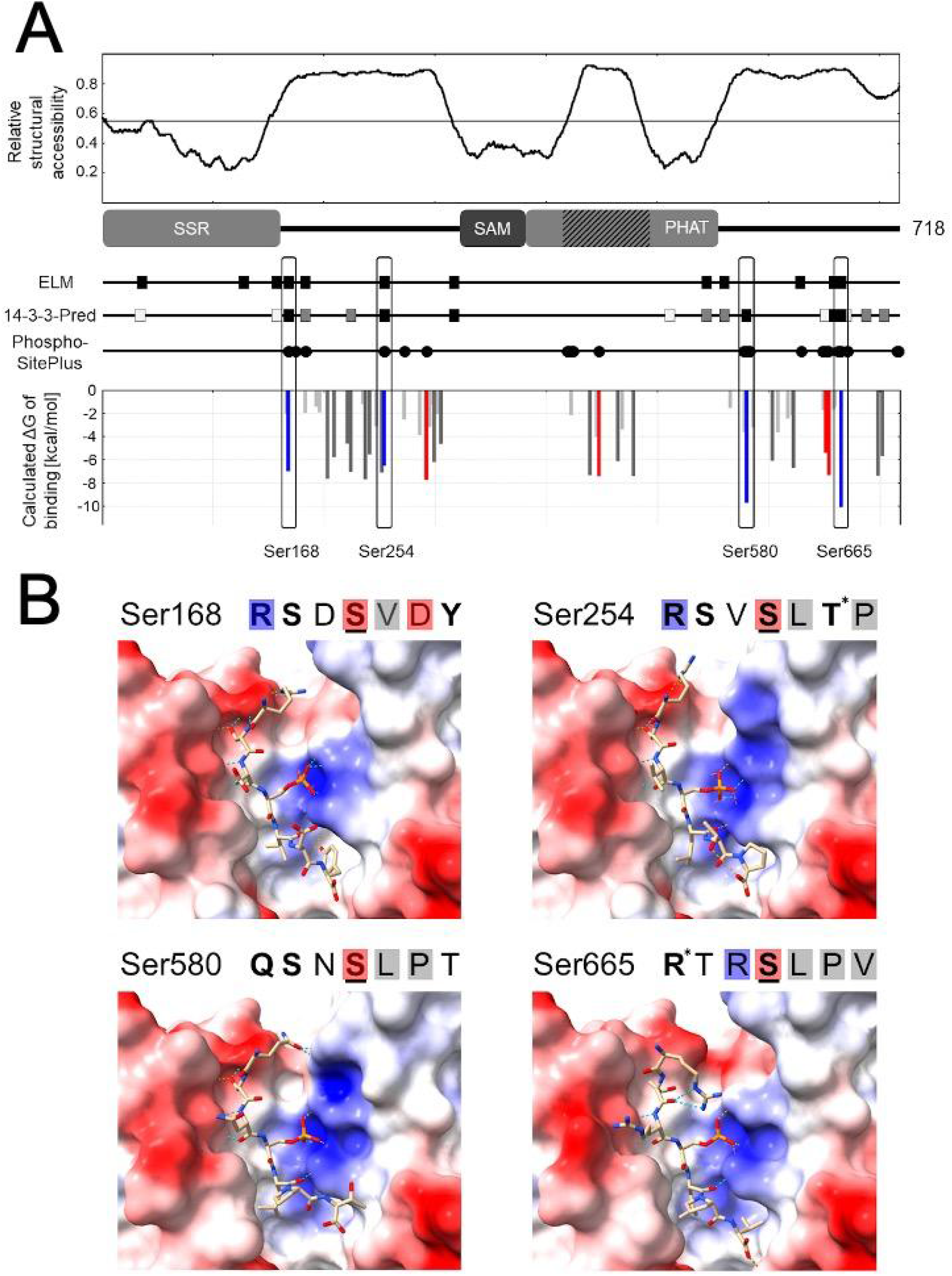
Computational identification of four high confidence 14-3-3 binding motifs in SMAUG1. (A) The top graph shows the relative structural accessibility calculated from the AlphaFold2 structural model. The lines below the domain diagram of SMAUG1 show candidate motif hits determined by ELM (black boxes) or 14-3-3-Pred (empty, grey or black boxes reflecting increasing reliability of the hit). Below the motif candidates, circles mark serine and threonine residues that have known phosphorylation sites in the PhosphoSitePlus database. The bottom part shows the estimated ΔG values calculated from predicted complex structures between Ser-centered SMAUG1 peptides and 14-3-3γ. Light grey bars mark interactions with poor estimated energies (ΔG > −4.1kcal/mol), dark gray bars mark peptides with no known phosphosites, red bars mark peptides that are highly unlikely to be able to bind to 14-3-3 domains in a protein context due to steric clashes, and blue bars denote high confidence binders. Bars mark the central serine residues of the tested peptides. (B) predicted complex structures of SMAUG1 14-3-3 motifs bound to 14-3-3γ. Peptide sequences are marked above each structure with the type of molecular interactions encoded in typesetting and colored boxes: the side chains of bold residues form hydrogen bonds with the domain, residues with blue and red boxes form strong electrostatic interactions with the domain via negative and positive charges of their side chains, residues with gray boxes form strong hydrophobic interactions with the domain, and residues with an asterisk form hydrogen bonds with the peptide backbone. Underlined residues mark the central phosphorylated serine residues. In the structures blue and red mark positively and negatively charged surface patches and dashed lines mark hydrogen bonds as assessed by ChimeraX [47].

**Figure 5:**
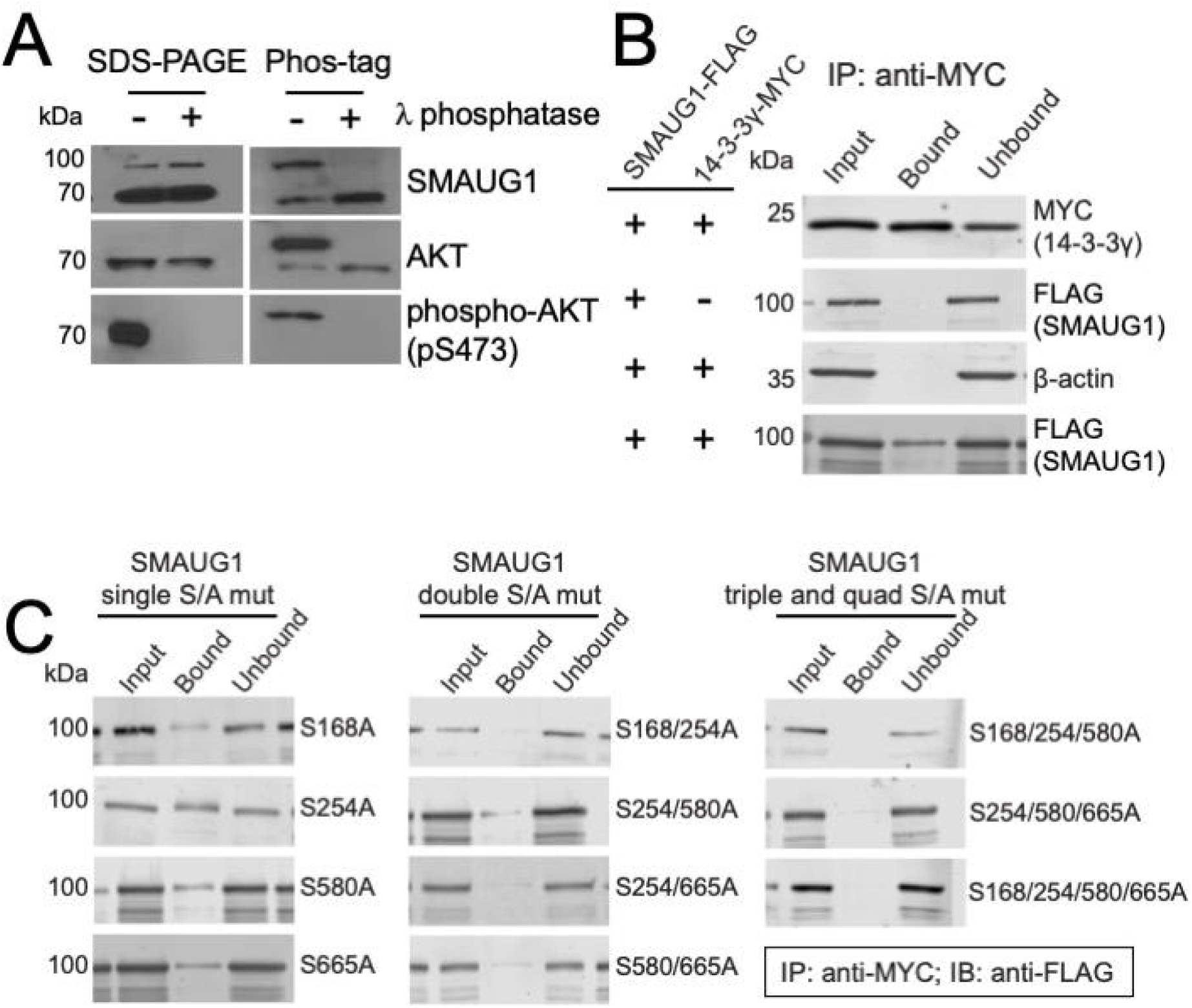
SMAUG1 is a phosphoprotein, and its 14-3-3 binding motifs control interactions with 14-3-3γ in a multivalent manner. (A) SMAUG1 phosphorylation status in HEK293 cells was determined using a Mn^+2^ Phos-tag SDS-PAGE gel. The phosphorylation status of SMAUG1 was determined through comparison of gel migration of lambda phosphatase-treated protein samples compared to untreated protein samples. Treated and untreated samples from whole-cell lysates were subjected to SDS-PAGE on 8% polyacrylamide gel containing 0 μM (no Mn2^+^–Phos-tag) or 25 μM Mn2^+^–Phos-tag, followed by immunoblotting with anti-SMAUG1, anti-AKT and anti-phospho-AKT Ser^473^. (B) Anti-myc co-immunoprecipitations with FLAG-tagged wild-type SMAUG1 and myc-tagged 14-3-3γ from co-transfected HEK293 cells. SMAUG1 is present in the bound fraction only when 14-3-3γ is co-expressed in the cells; β-actin was used as a negative control. (C) Anti-myc co-immunoprecipitations with FLAG-tagged SMAUG1 that has Ser to Ala mutations at positions 168, 254, 580 and 665 within four 14-3-3 binding motifs. Mutant FLAG-tagged versions of SMAUG1 were co-transfected with myc-tagged 14-3-3γ in HEK293 cells. Detection of SMAUG1 in the bound fraction decreases as the number of mutated 14-3-3 binding motifs increases. This indicates that 14-3-3γ interacts with SMAUG1 in a multivalent manner.

### Fluorescence microscopy of biomolecular condensates formed by SMAUG1 protein in cells

The ability of SMAUG1 deletion mutants (Figure 2) and the mutant with Ser to Ala changes in the four identified 14-3-3 binding motifs (quad S/A mutant; Figure 6 and Figure 7) to form condensates in cells was monitored by transient transfection of U-2 OS cells using FuGene (Promega; Figures 2 and 7) or electroporation (as before, Figure 6). For cells transiently transfected using FuGene, 2μg of plasmids SMAUG1-GFP, SMAUG1ΔIDR1, SMAUG1ΔSAM, SMAUG1ΔPLD (Figure 2) or 2μg of SMAUG1-GFP/quad S/A mutant SMAUG1-GFP was co-transfected with 1μg pCMV-14-3-3g-mCherry/pCMV-mCherry vector (Figure 7) according to the manufacturer’s protocol in 24-well plates. The transfection period ranged between 24-48 hours. Following removal of media, cells were washed 3 times with PBS and fixed with 4% PFA at room temperature for 15 minutes. After removal of PFA, cells were washed 3 times with PBS and cell nuclei were stained with DAPI. To determine the % granular cells for each transfection combination, fixed cells in PBS were imaged directly in 24-well dishes using an EVOS FL Auto Imaging System (20X objective). Counts of granular/non-granular cells were performed manually using the counting feature on the EVOS system. We observed clear differentiation between cells possessing condensates from those that had none or very few. Details of images collected, and number of cells counted are in Supplementary Tables S3 and S8.

**Figure 6:**
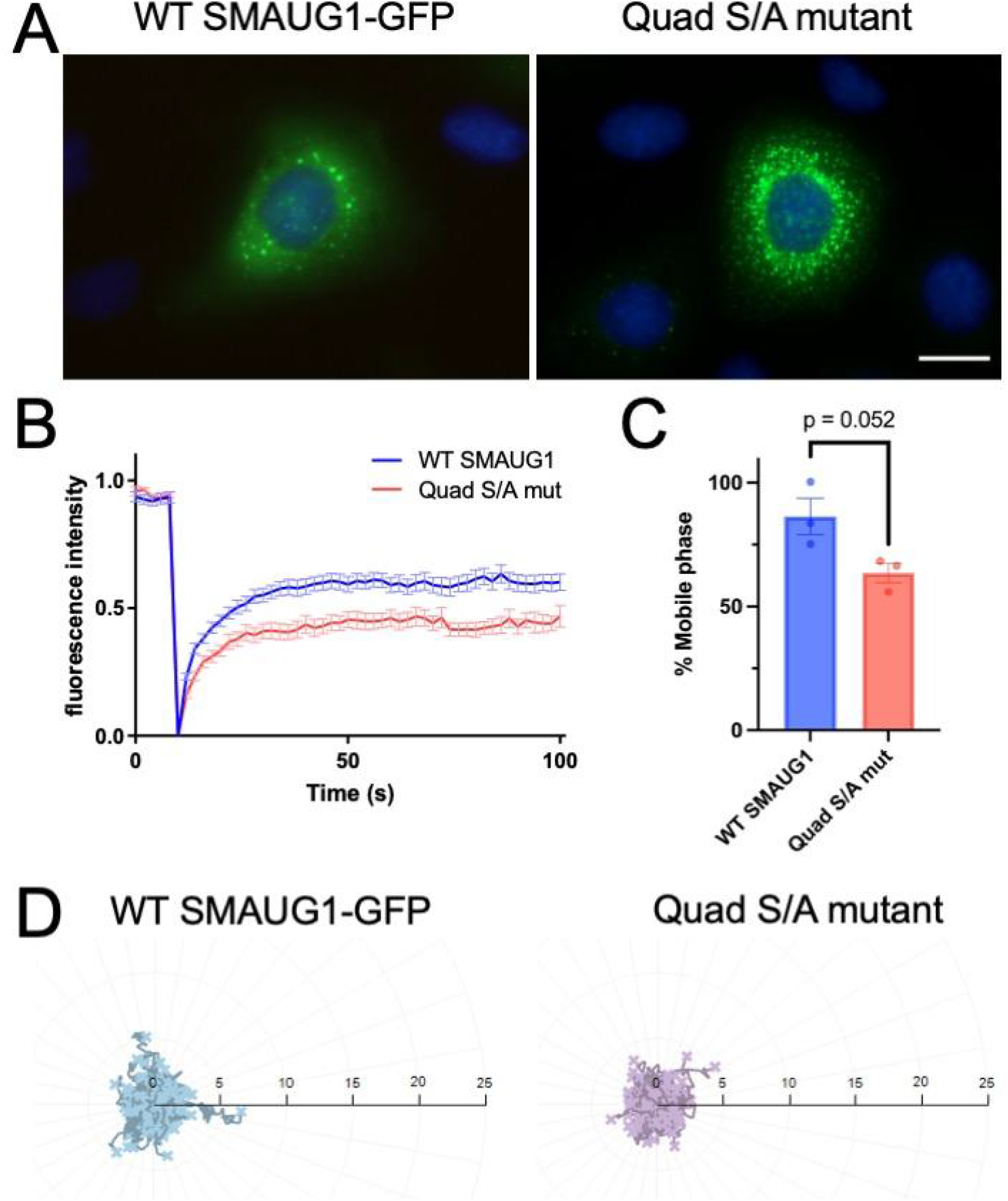
SMAUG1 condensates, with disrupted 14-3-3 interactions, show altered morphology and dynamics. (A) Fluorescence micrographs of U-2 OS cells expressing wild-type SMAUG1-GFP or mutant SMAUG1-GFP in which four, 14-3-3 binding motifs contained S to A mutations (quad S/A mutant) at positions 168, 254, 580 and 665. Cells transfected with quad S/A mutant SMAUG1-GFP show an increased number of condensates. Scale bar = 20μm (B) FRAP recovery curves of WT and mutant SMAUG1 protein following photobleaching. Images were recorded every 2 seconds; shown are pre-bleach, bleach (curves drop to 0), and post-bleach recovery for 100 seconds. Recovery of the photobleached region of interest was reduced with mutant SMAUG1 compared to the wild-type protein. For FRAP videos of the two constructs see Supplementary videos 1 and 2. (C) Determination of % mobile phase using FRAP analysis from three biological replicates. Average % mobile fraction is shown; bars are SEM. Means of WT and quad S/A mutant were compared with an unpaired, two-tailed t-test (p = 0.052) using GraphPad Prism 9. (D) Trajectory tracks of condensates formed by WT and quad S/A mutant SMAUG1-GFP protein created using FastTrack AI (MetaVi Labs). Scale bar is in μm. For videos of trajectory tracks, see Supplementary videos 3 and 4.

**Figure 7:**
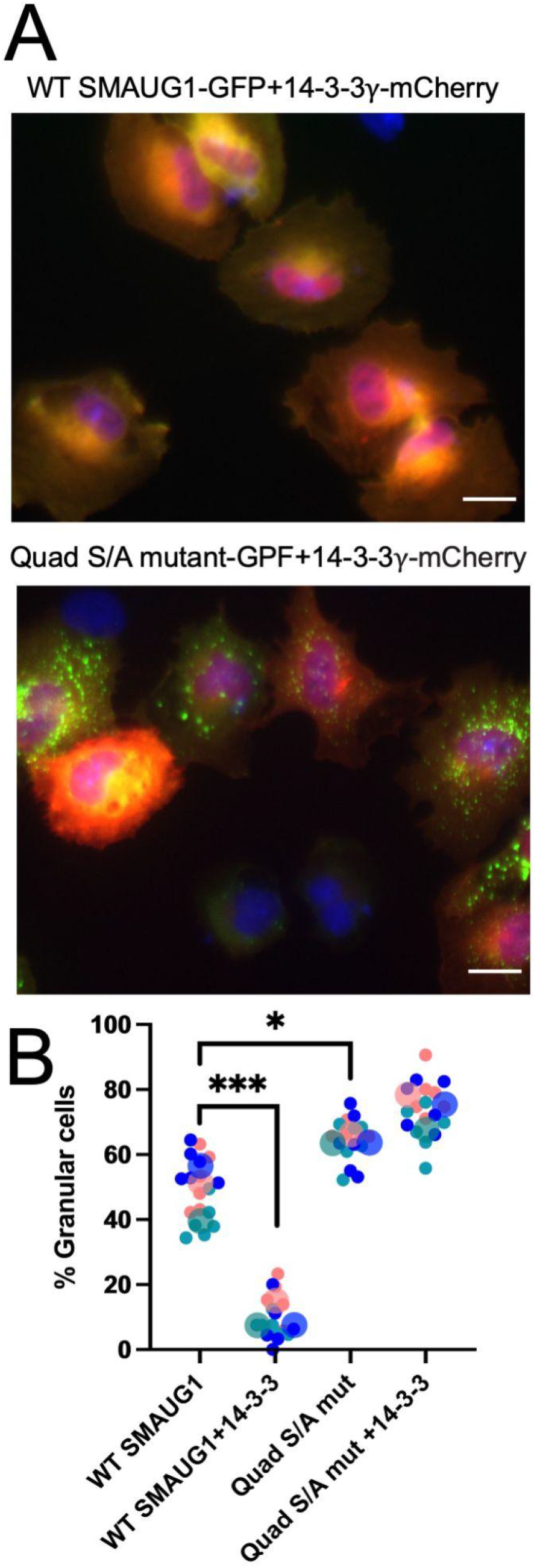
Co-expression of SMAUG1 and 14-3-3γ in cells leads to condensate dissolution. (A) Representative images of U-2 OS cells that were transiently co-transfected with either WT SMAUG1-GFP or Quad S/A mutant SMAUG1-GFP and 14-3-3γ-mCherry. WT SMAUG1 shows a significantly lower number of cells with condensates when 14-3-3γ is co-expressed. This effect is not observed when 14-3-3γ is co-expressed with the quad S/A SMAUG1 mutant. Images were collected at 60x magnification with the scale bar = 50μm. (B) Superplots [44] graph showing the percentage of granular cells when co-transfected with WT SMAUG1-GFP and 14-3-3γ-mCherry. Each color represents a biological replicate; the large circle is the average % granular cells of the technical replicates (small circles; n=5). Means of each group were compared with a one-way ANOVA using GraphPad Prism 9 using the Tukey’s test to correct for multiple comparisons, p<0.05 (*), p<0.001 (***).

## Results

### Human SMAUG1 drives the formation of liquid-like condensates in cells via its prion-like domain

Previous work has shown that SMAUG1 forms condensates in neurons and osteosarcoma (U-2 OS) cells [8,18,19]. Since SMAUG1 has been observed to partially overlap with markers of stress granules (SGs) in some cell types [18], we examined the localization of SMAUG1, fused at the C-terminus with enhanced green fluorescent protein (EGFP) (SMAUG1-GFP), following transient transfection into U-2 OS cells (Figure 1). We used indirect immunofluorescence to examine the localization of processing body (P-body) and SG markers in transfected cells, since both condensates are known to contain a high number of mRNAs, and the RNA-binding capacity of SMAUG1 makes them likely candidates for co-localization. P-bodies are continually present in the cell, however, SGs form only in response to cellular stress. To chemically induce SG assembly, transfected cells were treated with sodium arsenite (NaAsO_2_) for one hour, to induce oxidative stress, and returned to normal media [35].

We observed that SMAUG1-GFP co-localized with P-bodies and SGs by indirect immunofluorescence using antibodies directed against enhancer of mRNA-decapping protein 4 (EDC4), a known component of P-bodies [36,37] (Figure 1A top), and Ras-GTPase-activating binding protein 1 (G3BP1), a protein that nucleates SG assembly [38] (Figure 1A bottom). Interestingly, unlike previous reports that SMAUG1 is not present in P-bodies [18], we observed that it partially overlaps with P-bodies based on co-localization with EDC4. Importantly, while SMAUG1 appears to be present or near these known condensates (Figure 1A, white arrows), it is not uniformly distributed and also forms distinct condensates that are not positive for EDC4 or G3BP1 (Figure 1A, black arrows).

To test the biophysical characteristics of the observed SMAUG1 condensates, we carried out fluorescence recovery after photobleaching (FRAP) analysis on U-2 OS cells transiently expressing SMAUG1-GFP. While care must be taken in the interpretation of FRAP data [39], we used this technique to simply assess fluidity of SMAUG1 condensates. No previous studies reported FRAP analysis for SMAUG1 condensates; however, FRAP data on Vts1/Smaug, the yeast homolog, showed minimal recovery following bleaching, suggesting a high viscosity or partial aggregation [16]. Our FRAP data shows that SMAUG1 displays very fast recovery (5 and 10 seconds, post-bleach), typical of liquid-liquid phase-separated condensates (Figure 1B).

While our results clearly show that SMAUG1 resides in unique condensates, the protein can be a driver (“scaffold”), a regulator, or a client in the condensation process [40,41]. If SMAUG1 drives the condensation, there must be distinct regions in the SMAUG1 sequence that mediate the necessary molecular interactions. To test this, we designed three deletion constructs, abrogating known or predicted regions that drive condensation in other phase separation systems. The deleted region in ΔIDR (intrinsically disordered region, residues 237-311) is a long contiguous stretch of amino acids that are predicted to be disordered and to bind to structured protein partners, potentially through short linear motifs (SLiMs) based on IUPred2A and ANCHOR2 [26]. We also tested a deletion of the RNA-binding SAM domain (ΔSAM, residues 321-381), as many condensates are formed and maintained through RNA interactions [42]. Finally, many proteins are known to drive condensation through regions with low sequence complexity, resembling yeast prion domains [43]. These prion-like domains (PLDs) are typically highly flexible, showing a polymer-like behavior forming non-stoichiometric interactions driven by charged, polar and aromatic residues. The region deleted in the ΔPLD construct (residues 567-605) is a candidate PLD with a high number of positively charged and polar residues (18% and 23%, respectively), which exhibits the strongest propensity for being a PLD by PLAAC [27] (see supplementary Figure S1).

The condensation properties of the three GFP-tagged, SMAUG1-deletion constructs were tested by transfecting U-2 OS cells, along with wild-type SMAUG1 and a GFP control (Figure 2B). Both ΔSAM and ΔIDR SMAUG1 form condensates comparable to wild type SMAUG1; however, cells transfected with ΔPLD SMAUG1 are devoid of SMAUG1 condensates, indicating that the PLD region is necessary for condensate formation. To quantitate this effect, images of cells from three biological replicates, each with five technical replicates, were visually scored for % granularity (Figure 2C and Supplementary table S3). Our results indicate that the PLD region in SMAUG1 is responsible for driving condensate formation, while IDR (237-311) and the RNA-binding domain (SAM, 321-381) are dispensable.

### SMAUG1 interacts with members of the 14-3-3 protein family

Earlier studies have identified mouse Smaug (Samd4) as a regulator of mammalian target of rapamycin complex 1 (mTORC1) signaling; a single amino acid change that disrupts this regulation led to severe dysregulation of metabolism and energy balance [7]. As this regulation was found to be connected to mouse Smaug’s ability to interact with 14-3-3 proteins, directly or indirectly, we set out to test whether human SMAUG1 interacts with members of the 14-3-3 family. High-throughput studies already indicated interactions between human SMAUG1 and 14-3-3 proteins [45]. We carried out co-immunoprecipitation (co-IP) experiments to confirm the interactions *in vitro* (Figure 3).

Figure 3A shows that GFP-tagged human SMAUG1 pulls down endogenous 14-3-3 proteins from HEK293 cell lysates, as evidenced by identification with a pan-14-3-3 antibody, following immunoblotting. To understand whether this indicates an interaction with a single type of 14-3-3 protein or is a result of aspecific interactions with several, or all, members of the 14-3-3 family, we tested interactions with specific 14-3-3 proteins. Figure 3B shows that FLAG-tagged SMAUG1 pulls down both 14-3-3η and 14-3-3ζ, confirming that SMAUG1 can interact with more than one member of the 14-3-3 family.

Conversely, we also tested if 14-3-3 proteins can pull down endogenous SMAUG1 (see Figure 3C). For this, myc-tagged 14-3-3α/β, 14-3-3γ, 14-3-3η and 14-3-3ζ were overexpressed in HEK293 cells and immobilized with anti-myc magnetic beads. SMAUG1 was present in the immunoprecipitations for all 14-3-3 family members tested. This confirms that SMAUG1 does not exclusively bind a single member of the 14-3-3 family, and that the SMAUG1:14-3-3 interaction is not a result of the abnormally high levels of SMAUG1 due to overexpression.

While these results show that there is an interaction between SMAUG1 and various members of the 14-3-3 protein family, we cannot assess whether this is a direct physical interaction, or an indirect interaction mediated by some ancillary molecule(s). To rule out the involvement of ancillaries, we first tested whether the interaction depends on the presence of RNA, as SMAUG1 is known to interact with mRNAs [8,19]. Figure 3D shows that the SMAUG1:14-3-3 interactions form even in the absence of RNA following RNase treatment of cell lysates. This indicates that the interaction is RNA-independent and mediated by direct SMAUG1:14-3-3 contact or by additional protein(s).

### SMAUG1 contains four high-confidence 14-3-3 binding motif candidates

Our results in the previous section show that SMAUG1 interacts with members of the 14-3-3 protein family. However, it is not clear if this interaction is direct or is mediated by an ancillary protein. Thus, as the next step, we assessed the capacity of SMAUG1 to form direct physical contact with 14-3-3 domains, which recognize short linear motifs (SLiMs) in the sequences of their partner proteins [46]. As these SLiMs must fit into the glove-like structure of the 14-3-3 domain, they need to be in a structurally highly accessible/disordered protein region to be functional. 14-3-3-binding SLiMs are invariably centered around a phosphorylated serine (Ser) or threonine (Thr) residue fitting into a deep positively charged pocket inside the 14-3-3 domain. This pSer or pThr residue is flanked by other charged and hydrophobic residues that form additional contacts increasing specificity and binding strength.

We used two sequence-based SLiM identification tools, the Eukaryotic Linear Motif (ELM) resource [33], and the more specialized 14-3-3-Pred [34] to find candidates of 14-3-3 binding motifs. Figure 3 and Supplementary table S4 show that the two methods altogether predict 21 instances, which is improbably high. To arrive at a more realistic list of 14-3-3 motif candidates, we constructed a structure-based 14-3-3 SLiM identification pipeline. This pipeline considers three aspects of regions capable of 14-3-3 binding: 1) residing in a structurally accessible/disordered region, 2) sequence and structural compatibility with 14-3-3 binding, and 3) having a phosphorylatable residue in the middle. In the first step, we assessed structural accessibility of SMAUG1 calculated from the AlphaFold2 structural model (see Methods). Based on this [12], the regions connecting the domains, as well as the region inserted into the PHAT domain are disordered (Figure 3A, Supplementary table S5). In the second step, we tested every possible 7-mer peptide, with a serine in the middle, from the disordered regions of SMAUG1 for sequential and structural compatibility with 14-3-3 binding. For each peptide we built a predicted complex structure with human 14-3-3γ. We used an experimentally determined, high-quality complex structure from PDB as a template, introduced mutations to transform the peptide sequence to that of the SMAUG1 peptide being tested, and employed backbone and side-chain optimization to arrive at a predicted complex structure. Since 14-3-3 binding requires phosphorylation, we modeled each peptide with a pSer in the middle, regardless of the presence or absence of known phosphosites. This framework employs a combination of the FoldX and FlexPepDock methods to perform mutant generation and structure optimization, as both methods can handle modified residues, as opposed to AlphaFold2. As a result, for each User-centered SMAUG1 candidate peptide, we obtained the best complex structure, together with the estimated ΔG Gibbs free energy of the binding (see Methods). In the third step, we considered known phosphorylations for SMAUG1 from PhosphoSitePlus (Supplementary table S6).

Figure 3A and Supplementary table S7 shows the results of our structure-based 14-3-3 SLiM identification. Several SMAUG1 peptides yield a relatively high ΔG value that is not compatible with stable binding. Peptides without known phosphorylation sites can also be discarded, regardless of predicted binding energies. Finally, certain peptides that harbor known phosphorylation sites yield large negative (favorable) binding energies, however, looking at their predicted structures it is clear that they would have strong steric clashes with the 14-3-3 in a protein context, as their N- or C-termini point towards the domain surface and thus these binding modes are not accessible for a full-length SMAUG1 protein (for all modeled complex structures see Supplementary Dataset 1).

After discarding all incompatible hits, four high-confidence SLiM candidates are left, centered around Ser168, Ser254, Ser580 and Ser665. All four of these are also among the candidate list of 14-3-3-Pred and three are also identified by ELM. To assess if the binding modes for these SLiM candidates are indeed realistic and do resemble the known 14-3-3:peptide complex structures, Figure 3B shows the predicted bound structures. For each four, the peptide binds in the correct orientation and forms various molecular contacts with the domain that are known to occur in 14-3-3 binding. Ser168, Ser254 and Ser665 all have the common Arg residue in the −3 position, each peptide forms hydrophobic contacts C-terminal of the pSer residue, and all four peptides form a high number of hydrogen bonds with residues of the domain.

### SMAUG1 contains four phospho-dependent 14-3-3 binding motifs providing multivalent interactions

Following on from our computational identification of four high-confidence 14-3-3 binding motifs in SMAUG1, we next wanted to investigate the contribution of each of these motifs to the interaction with 14-3-3γ *in vitro*. Since the four 14-3-3-binding SLiMs identified are centered around a presumably phosphorylated serine at positions 168, 254, 580 and 665, we first examined the phosphorylation status of SMAUG1 in HEK293 cell lysates. We prepared whole cell lysates, with the addition of phosphatase inhibitor, and resolved the proteins on SDS-PAGE or phos-tag gels, followed by immunoblotting with anti-SMAUG1 antibody. Phos-tag gel electrophoresis allows for the simultaneous detection of multiple phospho-sites in a protein using Mn^+2^ or Zn^+2^ ions that bind to the phosphate groups [48,49]. The interaction with manganese or zinc cations shifts the mobility of phosphoproteins in the gel system, and the change in mobility can be easily visualized. Figure 5A shows that SMAUG1 is phosphorylated (higher mobility band in the phos-tag gel); however, to confirm that the change in mobility was due to phosphorylation, we treated half of the lysate with lambda protein phosphatase. With the addition of phosphatase, the higher mobility SMAUG1 band disappears, and what is detected is the unphosphorylated protein. As a control we also monitored the phosphorylation of AKT (protein kinase B) using a phospho-specific antibody against pSer473 (Figure 5A).

After we confirmed that SMAUG1 is phosphorylated, we next transfected FLAG-tagged SMAUG1 and myc-tagged 14-3-3γ into HEK cells to test their interaction by immunoprecipitation with anti-myc beads, followed by immunoblotting. Figure 5B shows that myc-tagged 14-3-3γ is enriched in the bound fraction (top panel) and can pulldown FLAG-tagged SMAUG1 (bottom panel). Without 14-3-3γ, SMAUG1 cannot be precipitated.

Following the confirmation of the interaction of wild-type SMAUG1 with 14-3-3γ by co-immunoprecipitation, we created a series of serine to alanine mutations in SMAUG1 at positions 168, 254, 580 and 665 (S168A, S254A, S580A and S665A) using site-directed mutagenesis of the FLAG-tagged SMAUG1 construct. In addition to single amino acid changes, we also created double, triple, and quad mutant combinations to test the valency of SMAUG1’s interaction with 14-3-3γ. We then tested the ability of FLAG-tagged SMAUG1 mutants to interact with myc-tagged 14-3-3γ in HEK293 cells. Figure 5C indicates that single site mutations had only a negligible effect on SMAUG1 binding; whereas the double mutants showed only a residual amount of SMAUG1 present in the bound fraction (second panel). Mutating three or all four 14-3-3 binding motifs in SMAUG1 appeared to eliminate the interaction with 14-3-3γ as evidenced by the lack of SMAUG1 in the bound fractions (third panel). These data indicate that the phosphoserine residues within the four 14-3-3 binding motifs identified in SMAUG1 are critical for the protein’s interaction with 14-3-3 proteins in a multivalent manner.

### Disrupted 14-3-3 interactions in SMAUG1 alter condensate morphology and dynamics

Having identified four 14-3-3 binding motifs and confirmed that they control SMAUG1 interactions with the 14-3-3 proteins in a phospho-dependent manner, we next wanted to see if disruption of these interactions influenced SMAUG1’s ability to form condensates. First, we looked at overall differences in SMAUG1 condensate morphology using fluorescence microscopy (Figure 6A). Condensates were observed in U-2 OS cells at 24-hours post-transfection with wild-type SMAUG1-GFP and mutant SMAUG1-GFP, which contained the four serine to alanine mutations S168A, S254A, S580A and S665A (quad S/A mutant). In comparison to the wild-type protein, Figure 6A shows that cells containing SMAUG1 with disrupted 14-3-3 binding motifs show an increased number of condensates. This indicates that the mutant protein still forms condensates; however, disabling the interactions with 14-3-3 protein prevents condensate dissolution. The observed result is that the cell becomes packed with condensates that cannot dissipate.

We tested the dynamics and biophysical properties of wild-type and mutant SMAUG1 condensates in live cells using FRAP (see Supplementary Materials, videos 1 and 2). Condensates in U-2 OS cells containing wild-type or mutant GFP-SMAUG1 were photobleached and monitored for fluorescence recovery. Comparison of recovery curves revealed that quad S/A mutant SMAUG1 displayed a lower rate of exchange with the cytoplasm than the wild-type protein (Figure 6B, left). Furthermore, the mobile fraction of WT SMAUG1 was 86.4% and 63.5% for the mutant protein (Figure 6B, right). Still our FRAP data shows that 14-3-3-binding incompetent quad-mutant SMAUG1 displays very fast recovery, indicating that disabling the interaction with 14-3-3 proteins does not fundamentally change the liquid properties of SMAUG1 condensates.

Next, we wanted to examine the movement of wild-type and mutant SMAUG1 condensates within an individual cell. Using the FRAP videos that we collected, non-photobleached condensates were tracked using a modification of the chemotaxis application within FastTrack AI (MetaVi Labs). By tracing the path of each condensate, we obtained trajectory track videos of wild-type and mutant GFP-SMAUG1 (see Supplementary Material, videos 3 and 4). To capture static images that represent the movement of the SMAUG1 condensates, the trajectory of each condensate was recorded and plotted. Figure 5D shows the trajectory plots of wild-type and mutant SMAUG1 condensates in the transfected U-2 OS cells. Interestingly, wild-type and mutant condensates display slightly altered mobility; therefore, the disruption of 14-3-3 binding motifs likely affects other protein interactions that are important for condensate movement and/or anchoring, in addition to regulating SMAUG1 condensate dissolution.

### SMAUG1:14-3-3 interactions modulate SMAUG1 condensation in cells

While our fluorescence micrographs, FRAP and condensate tracking of wild-type and mutant GFP-tagged SMAUG1 showed noticeable differences due to the disruption of 14-3-3 binding motifs, the resultant effects relied on endogenous levels of 14-3-3 proteins in cells. To have direct observations that monitor the effect of 14-3-3 protein interactions with SMAUG1, we co-transfected U-2 OS cells with wild-type GFP-tagged SMAUG1 (SMAUG1-GFP) or the quad mutant (S168/254/580/665A, used in previous experiments) together with mCherry-tagged 14-3-3γ (14-3-3γ-mCherry). In Figure 7A, 14-3-3γ shows high diffuse expression throughout the cytoplasm, regardless of the mutation status of SMAUG1-GFP in co-transfected cells. SMAUG1 distribution; however, is strikingly different, depending on whether the 14-3-3 binding motifs are intact (wild-type SMAUG1-GFP, Fig 7A top) or have been mutated (quad S/A mutant SMAUG1-GFP, Fig 7A bottom).

In cells co-expressing wild-type SMAUG1 and 14-3-3γ, SMAUG1 displays similar diffuse cytoplasmic localization to 14-3-3γ, and the cells are largely devoid of condensates. In contrast, the quad mutant SMAUG1, which cannot bind to 14-3-3 proteins through any of its four motifs, shows a high degree of condensation, despite the presence of overexpressed 14-3-3γ. These results further demonstrate that 14-3-3 is a strong negative regulator of SMAUG1 condensation, as interactions between the two molecules lead to the dissolution of SMAUG1-driven condensates.

Apart from demonstrating the strong negative regulatory effect of 14-3-3 in SMAUG1 condensation (Figure 7A), we also collected a large amount of quantitative data on the number of co-transfected cells that possess SMAUG1 condensates (Figure 7B; Supplementary table S8). As our previous data shows, 14-3-3 is highly unlikely to inhibit SMAUG1 condensation, but instead, it aids the dissolution of condensates. Hence, even in the presence of exogenous 14-3-3, some cells might be able to display SMAUG1 condensates due to the high relative concentration of SMAUG1 vs 14-3-3γ, the presence of a third molecule that modulates the condensation process or the inhibitory role of 14-3-3γ binding, or some other, unknown mechanism. Figure 7B shows the average percentage of cells that were categorized as granular (see Materials and Methods), for four different combinations of constructs used for transfections. In the first case, we co-transfected U-2 OS cells with wild-type SMAUG1-GFP and mCherry vector; SMAUG1-GFP formed condensates in most observed cells. In the second case, we co-transfected the cells with both wild-type SMAUG1-GFP and 14-3-3γ-mCherry, and the average fraction of granular cells dropped to about 10%. In the third case, we co-transfected the cells with quad mutant SMAUG1-GFP, which is 14-3-3 binding incompetent, and mCherry vector. Expression of mutant SMAUG1-GFP resulted in the formation of condensates in about 65% of the observed cells. This fraction is higher than that observed in cells transfected with wild-type SMAUG1 with weak significance, due to the abolishment of the negative regulation by endogenous 14-3-3. Since the abundance of overexpressed mutant SMAUG1 is likely much higher than that of 14-3-3γ, this increase is noticeable but not dramatic. Unexpectedly, in the fourth case where we co-transfected the cells with the quad mutant SMAUG1-GFP and 14-3-3γ-mCherry, we observed an even higher fraction of granular cells of about 75%, noticeably higher than in the wild-type case. Since in the third and fourth cases SMAUG1 is presumed to be incapable of binding to 14-3-3γ, we cannot easily explain this increase in the context of direct interactions between the two proteins. Indeed, while SMAUG1:14-3-3 phospho-dependent interactions are partially controlling SMAUG1 condensation, there are likely more complex regulatory mechanisms at play that will require further experiments to decipher.

## Discussion

The results we present in this paper provide mechanistic insights into how SMAUG1 condensation is initiated and modulated through interactions regulated by phosphorylation. The ability of SMAUG1 to reside in a condensed phase has been long recognized as one of its well-conserved functions [18,19]. Our work recapitulates this behavior in a cellular environment, and we also show that the resulting condensates are formed via liquid-liquid phase separation, as they are highly dynamic (Figure 1). The driver of SMAUG1 condensation is the C-terminal 567-605 region (Figure 2) that bears a strong resemblance to prion-like domains (PLDs) that can form non-stoichiometric assemblies via homotypic interactions [21]. Despite this resemblance, SMAUG1 567-605 seems to be unique in the sense that methods specifically trained to recognize PLDs [27] fail to give a high confidence prediction on SMAUG1 (see Supplementary Figure S1), and methods trained to recognize disordered interaction sites mediating stoichiometric protein-protein interactions [26] give a high score (Figure 2A), indicating that SMAUG1 567-605 might have a wider interactional repertoire than most PLDs.

Apart from residing in condensates, the other well-conserved function of SMAUG1 is its RNA-binding ability. While RNA binding is critical for initiating condensation for many phase separation drivers (such as nucleophosmin [50], fragile X mental retardation protein (FMRP) [51] and non-POU domain-containing octamer-binding protein (NONO) [52]), SMAUG1 is an exception, as its condensation seems to be independent of RNA binding (Figure 2). Instead, it is likely the reverse scenario, where the RNA-binding and processing capabilities of SMAUG1 are regulated by whether it is in a condensed or a dispersed phase. It would be an exciting future exploration to understand the exact relationship between the condensation and RNA-binding functions of SMAUG1, especially so, because SMAUG1 not only forms its own condensates, but can contribute to forming other condensates, such as stress granules and P-bodies (Figure 1). The role of RNA is well-established in both membraneless organelles [37,53], and SMAUG1 might be able to alter their RNA composition and functionalities. Both stress granules and P-bodies can form without SMAUG1, which suggests that, in addition to its LLPS driver function, SMAUG1 can behave as an LLPS regulator or client in other molecular settings, indicating a multifaceted role.

In the context we examined, SMAUG1 is an LLPS driver. Additionally, we identified a strong regulator of SMAUG1 condensation, namely the 14-3-3 protein family. Using a combination of sequence and structure-based computational methods, we identified four separate, phospho-regulated 14-3-3 recognition motifs in SMAUG1 (Figure 4) and verified these interactions using both co-IP assays using cell lysates and in cell measurements (Figures 3 and 5–7). The interaction with 14-3-3 proteins is a strong negative regulator of SMAUG1-mediated LLPS, as these interactions inhibit the formation or induce the dissolution of condensates. Judging by the co-IP experiments, the four motifs are likely acting cooperatively, as deletion of a single motif or two motifs seem to have a lesser effect than would be expected from linear models. However, the precise elucidation of this cooperativity will require fully quantitative assays with purified proteins and will need to be addressed in follow-up experiments.

While the interactions between SMAUG1 and 14-3-3 proteins could be predicted using computational methods (Figure 4), their negative regulatory effect on condensation is non-trivial. 14-3-3 proteins are dimers and are capable of binding two individual motifs. Thus, it is possible that a 14-3-3 dimer can bind two different copies of SMAUG1, increasing the valency, and forming larger molecular networks. These properties are known to aid condensation [54], and therefore, seeing 14-3-3 proteins exerting a positive regulatory role would not have been surprising. Our measurements showing a strong negative regulatory role, however, indicate a different dominant interaction pattern. A single 14-3-3 dimer binding to two motifs on the same SMAUG1 protein would in fact noticeably decrease the flexibility and overall degrees of freedom of SMAUG1, reducing the LLPS tendency. While our data clearly show that the result of the interaction is indeed the suppression of condensation, it is unclear how 14-3-3 dimers would be able to selectively prefer binding a single copy of SMAUG1 instead of linking two different copies. Exploring the interaction patterns between 14-3-3 and SMAUG1 molecules would enable us to address the role of multivalency in LLPS regulators, an aspect that is very poorly understood. These future experiments, however, would need to have excellent spatio-temporal resolution, as motif-mediated interactions tend to be transient and can typically form and break quickly.

While we focused on the role of 14-3-3 proteins, our results indicate that the regulation of SMAUG1 condensation is more complex than what we can directly measure. The major trends in our experiments clearly show the negative regulatory effect of 14-3-3, however, they do not explain a few phenomena we observed. The abundance of 14-3-3 proteins not only affects the number of condensates we see, but it also affects the number of granular vs non-granular cells. In addition, not all SMAUG1 expressing cells form condensates, even though 14-3-3 is not overexpressed in them. This might be explained by a few possible effects: 1) other negative regulators might be at play, 2) SMAUG1 can only drive condensation together with some other molecule that is missing from these cells, or 3) the expression of 14-3-3 in these cells is naturally higher due to some unknown mechanism. To explore these questions, we would need to follow up with measurements using purified proteins to have precise control over the molecules present in the experiment. These would be exceptionally valuable, as 14-3-3 regulation seems to be coupled to other, unknown regulatory mechanisms. This coupling is indicated by our data showing how adding 14-3-3 to a binding-deficient mutant of SMAUG1 has a slight positive effect on condensation (Figure 7). In addition, mutating the key phosphorylatable residues in the 14-3-3 motifs not only switches off the negative regulatory mechanism, but also changes the dynamics of SMAUG1 condensates (Figure 6).

SMAUG1 consists of 718 residues, and structural predictions show that roughly 400 of these are inside intrinsically disordered regions (IDRs). Roughly 10% of these residues are covered by the identified condensation-driving PLD region (Figure 2), and the four identified 14-3-3 motifs cover around an additional 5% - however, we still do not know what the rest of the disordered residues do. IDRs tend to have a high number of various interaction sites, including motifs, as well as modification sites, encoding functions in a very compact way [15,55,56]. This indicates that we are currently just scratching the surface of the SMAUG1 interactome, and thus, regulation and functionality. Mapping these interactions and understanding how they regulate SMAUG1 condensation and localization is of key importance to understand SMAUG1 function. Our current findings not only move us closer to a more complete understanding of SMAUG1 biology, but they have wider implications as well. While several motif-mediated interactions were described that help drive biological condensation [57–59], to our knowledge this is the first instance of 14-3-3 motifs shown to directly regulate LLPS. Previous studies have established a connection between the localization and function of the LLPS driver, nonphototropic hypocotyl 3 (NPH3) and its 14-3-3 binding capacity [60]. 14-3-3 binding was also shown to regulate whether tristetraprolin (TTP) resides in stress granules [61], thus 14-3-3 serves as a regulator of an LLPS client or co-scaffold, as stress granule assembly does not depend on the presence of TTP. Even without direct examples of 14-3-3 governing condensation, such as for SMAUG1, 14-3-3 proteins have been hypothesized to potentially regulate several condensation events [62]. Since 14-3-3 proteins are highly abundant in cells in virtually all human tissues [63], they are prime candidates to be general regulators of LLPS. The methods and findings we presented here can hopefully serve as both a motivation and template to uncover new functional modules in SMAUG1 and to investigate the wider role of 14-3-3 proteins in biological condensation.

## Abbreviations

SAM: sterile alpha motif
SAMD4A: SAM domain-containing protein 4A
PHAT: pseudo-HEAT analogous topology
SSR: SMAUG similarity region
PLD: prion-like domain
IDR: intrinsically disordered region
LLPS: liquid-liquid phase separation
SG: stress granule
P-body: processing body
AMPK: AMP-activated protein kinase
mTOR: mammalian target of rapamycin
EDC4: enhancer of mRNA-decapping protein 4 - UniProt
G3BP1: Ras-GTPase-activating binding protein 1
GFP/EGFP: green fluorescent protein/enhanced green fluorescent protein
PBS: phosphate buffered saline
FRAP: fluorescence recovery after photobleaching
ANOVA: analysis of variance
DSSP: define secondary structure of proteins
SLiM: short linear motif
ELM: Eukaryotic Linear Motif database

## Supplementary material

### Supplementary figures

**Figure S1:**
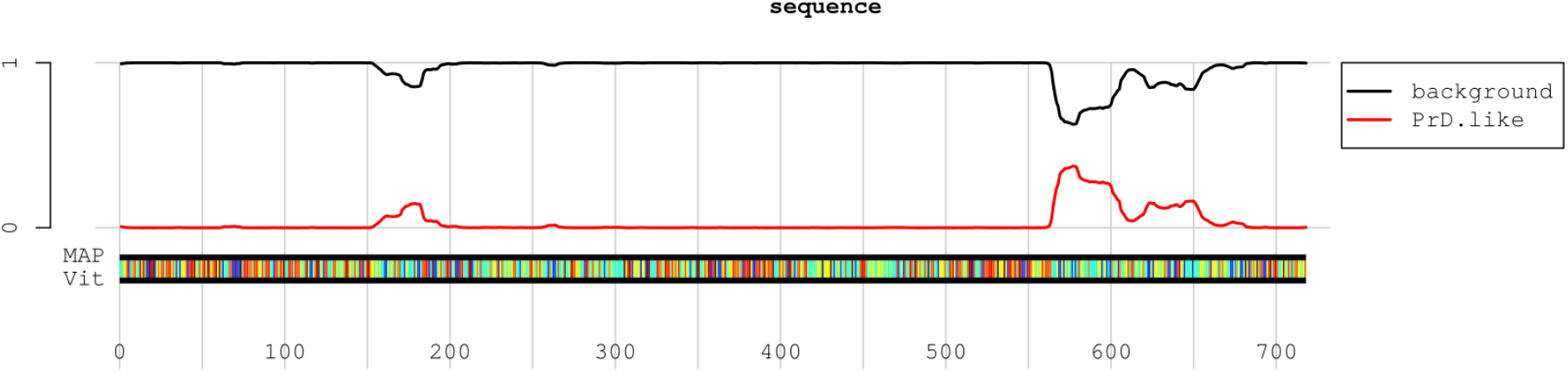
Prion-like domain prediction for human SMAUG1 using PLAAC. The red line represents the predicted tendency to form a prion-like domain.

### Supplementary tables

**Table S1: Sequences of gene blocks, DNA oligonucleotides, and mutagenic primers Table S2: Primary FRAP analysis measurements of WT/mutant SMAUG-GFP**

**Table S3: Granular cell counts from images of cell transfected with SMAUG1 deletion constructs**

**Table S4: SMAUG1 14-3-3 binding motif candidates identified from the sequence.** Numbers in the 14-3-3pred column show the number of sub-methods that give a positive prediction at that position.

**Table S5: Relative solvent accessible surface areas for residues in human SMAUG1.** Values are calculated based on AlphaFold2 structure and are averaged in a sliding window of ±15 residues.

**Table S6: Phosphorylation sites for human SMAUG1 in PhosphoSitePlus**

**Table S7: Candidate 14-3-3 interacting peptides from the disordered regions of human SMAUG1.** Light gray color marks interactions with poor estimated energies (ΔG > −4.1kcal/mol), dark gray marks peptides with no known phosphosites, red marks peptides that are highly unlikely to be able to bind to 14-3-3 domains in a protein context due to steric clashes, and blue denotes high confidence binders.

**Table S8: Granular cell counts from images of U-2 OS cells co-transfected with SMAUG1-GFP and 14-3-3gamma-mCherry**

### Supplementary datasets

**Supplementary dataset 1:** Predicted complex structures of candidate 14-3-3 binding peptides from the disordered regions of human SMAUG1 and human 14-3-3γ - <u>link</u>

### Supplementary videos

**Video 1:** Wild-type SMAUG1-GFP FRAP - <u>link</u>

**Video 2:** Quad S/A mutant SMAUG1-GFP FRAP - <u>link</u>

**Video 3:** Wild-type SMAUG1-GFP condensate tracking - <u>link</u>

**Video 4:** Quad S/A mutant SMAUG1-GFP condensate tracking - <u>link</u>

## Acknowledgements

We particularly would like to thank Nancy Kedersha for insightful discussions and help throughout the project. We are grateful to our School of Biochemistry and Cell Biology colleagues, Mary McCaffrey, Tom Moore and Paul Young, for valuable reagents and the use of equipment that greatly facilitated the project. We would like to thank Katja Burk and Cora O’Neill for proofreading the manuscript.

This work would not have been possible without the contributions of many undergraduate students including Enrica Antolini, Aparna Chandrasekaran, Niamh Daly, Emer Murphy, Sanjana Mathews and Cliodhna Wallace. Lastly, we thank members of the Dean, McCarthy, Moore and Young research groups for ongoing support and a collaborative working environment.

## Authors contributions

**John Fehilly:** Writing - original draft; Investigation; Formal analysis; Validation; Visualization

**Olivia Carey:** Writing - original draft; Investigation; Formal analysis; Validation; Visualization

**Eoghan Thomas O’Leary:** Writing - original draft; Formal analysis; Investigation; Visualization

**Stephen O’Shea:** Writing - original draft; Investigation; Visualization

**Klaudia Juda:** Writing - original draft; Investigation

**Rahel Fitzel:** Writing - original draft; Investigation

**Pooja Selvaraj:** Writing - original draft; Investigation

**Andrew J. Lindsay:** Writing - original draft; Investigation; Formal analysis; Writing - review & editing

**Bálint Mészáros:** Conceptualization; Data curation; Formal analysis; Methodology; Software; Visualization; Writing - original draft; Writing - review & editing

**Kellie Dean:** Conceptualization; Data curation; Formal analysis; Funding acquisition; Investigation; Methodology; Project administration; Supervision; Validation; Visualization; Writing - original draft; Writing - review & editing

